## Supplementary materials for "From sequence to molecules: Feature sequence-based genome mining uncovers the hidden diversity of bacterial siderophore pathways"

Pyoverdines analysed for the bioinformatic pipeline verification

| **Pyoverdine ID** | **MS / MS profiles available from** | |
|---|---|---|
| | **Rehm 2022 ABC** https://doi.org/10.1007/s00216-022-03907-w | **Gu 2024 eLife** https://doi.org/10.7554/eLife.96719.1 |
| 3A06 | 3A06 | x |
| 3B19 | 3B19 | x |
| 3G07 | 3G07 | x |
| s3b09 | s3b09 | x |
| s3c13 | s3c13 | x |
| s3e20 | s3e20 | x |
| 3A13 | x | 3A13 |
| 3A18 | x | 3A18 |
| 3B09 | x | 3B09 |
| 3C06 | x | 3C06 |
| 3C14 | x | 3C14 |
| 3D03 | x | 3D03 |
| 3E13 | x | 3E13 |
| s3a07 | x | s3a07 |
| s3a19 | x | s3a19 |
| s3b02 | x | s3b02 |
| s3d06 | x | s3d06 |
| s3e01 | x | s3e01 |
| s3e11 | x | s3e11 |
| s3e15 | x | s3e15 |

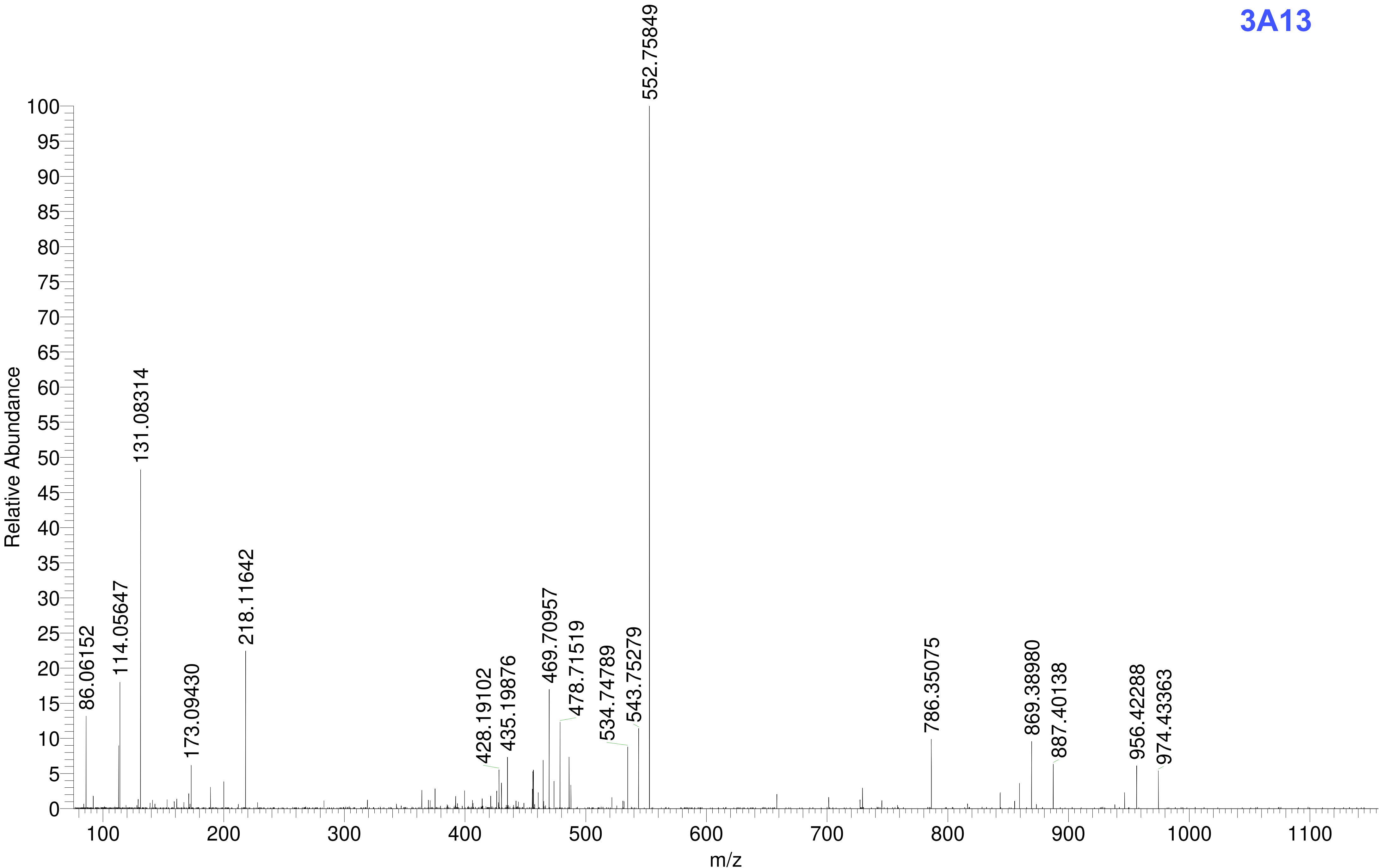

3A13
Relative Abundance
m/z
552.75849
131.08314
218.11642
114.05647
86.06152
173.09430
428.19102
435.19876
469.70957
478.71519
534.74789
543.75279
786.35075
869.38980
887.40138
956.42288
974.43363

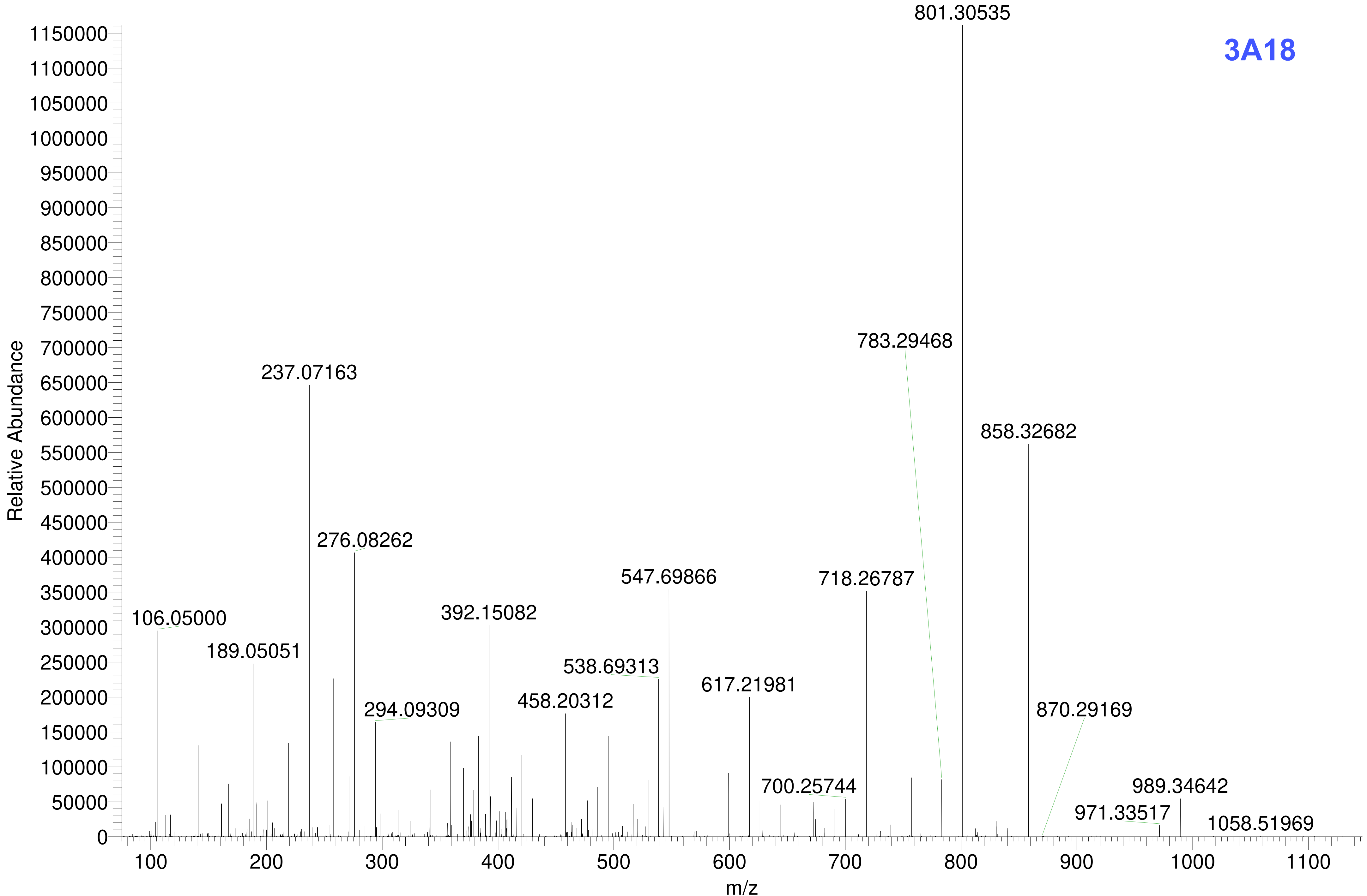
3A18
801.30535
783.29468
858.32682
237.07163
276.08262
547.69866
718.26787
392.15082
106.05000
189.05051
538.69313
617.21981
294.09309
458.20312
870.29169
700.25744
989.34642
971.33517
1058.51969
Relative Abundance
m/z

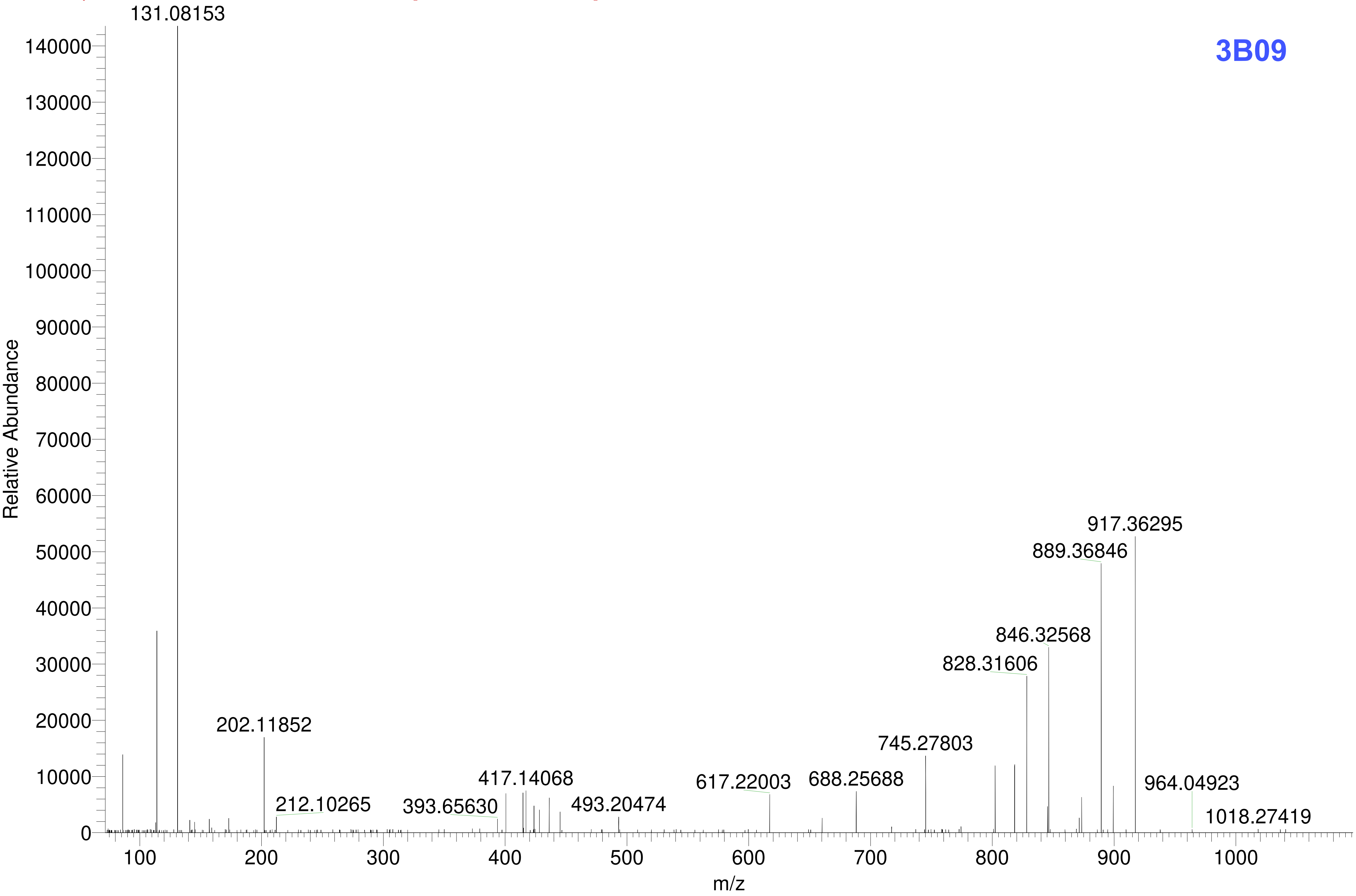

3B09
131.08153
202.11852
212.10265
393.65630
417.14068
493.20474
617.22003
688.25688
745.27803
828.31606
846.32568
889.36846
917.36295
964.04923
1018.27419
Relative Abundance
m/z

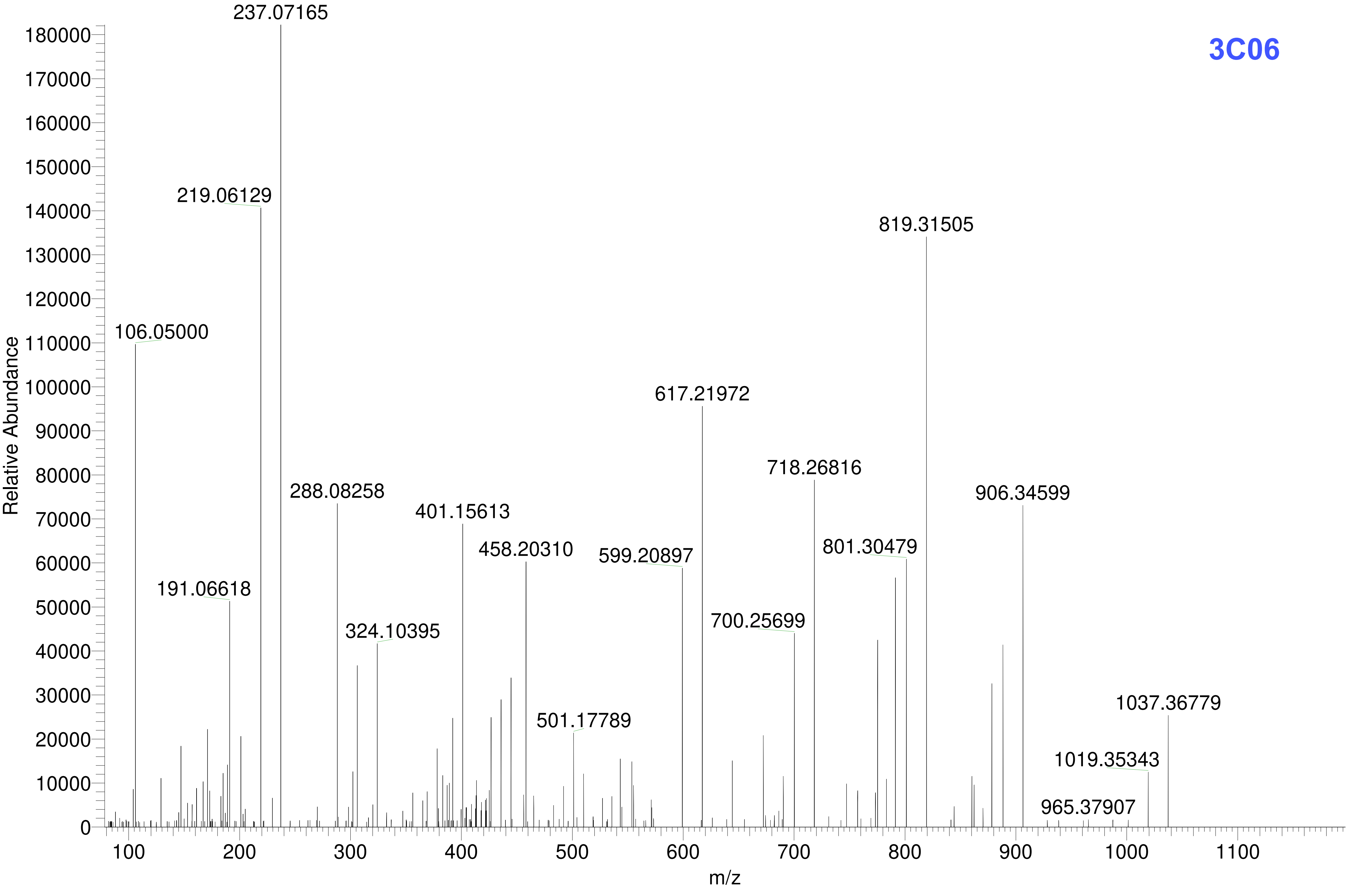
3C06
Relative Abundance
m/z
237.07165
219.06129
106.05000
819.31505
617.21972
718.26816
906.34599
288.08258
401.15613
458.20310
599.20897
801.30479
191.06618
324.10395
700.25699
1037.36779
501.17789
1019.35343
965.37907
180000
170000
160000
150000
140000
130000
120000
110000
100000
90000
80000
70000
60000
50000
40000
30000
20000
10000
0
100
200
300
400
500
600
700
800
900
1000
1100

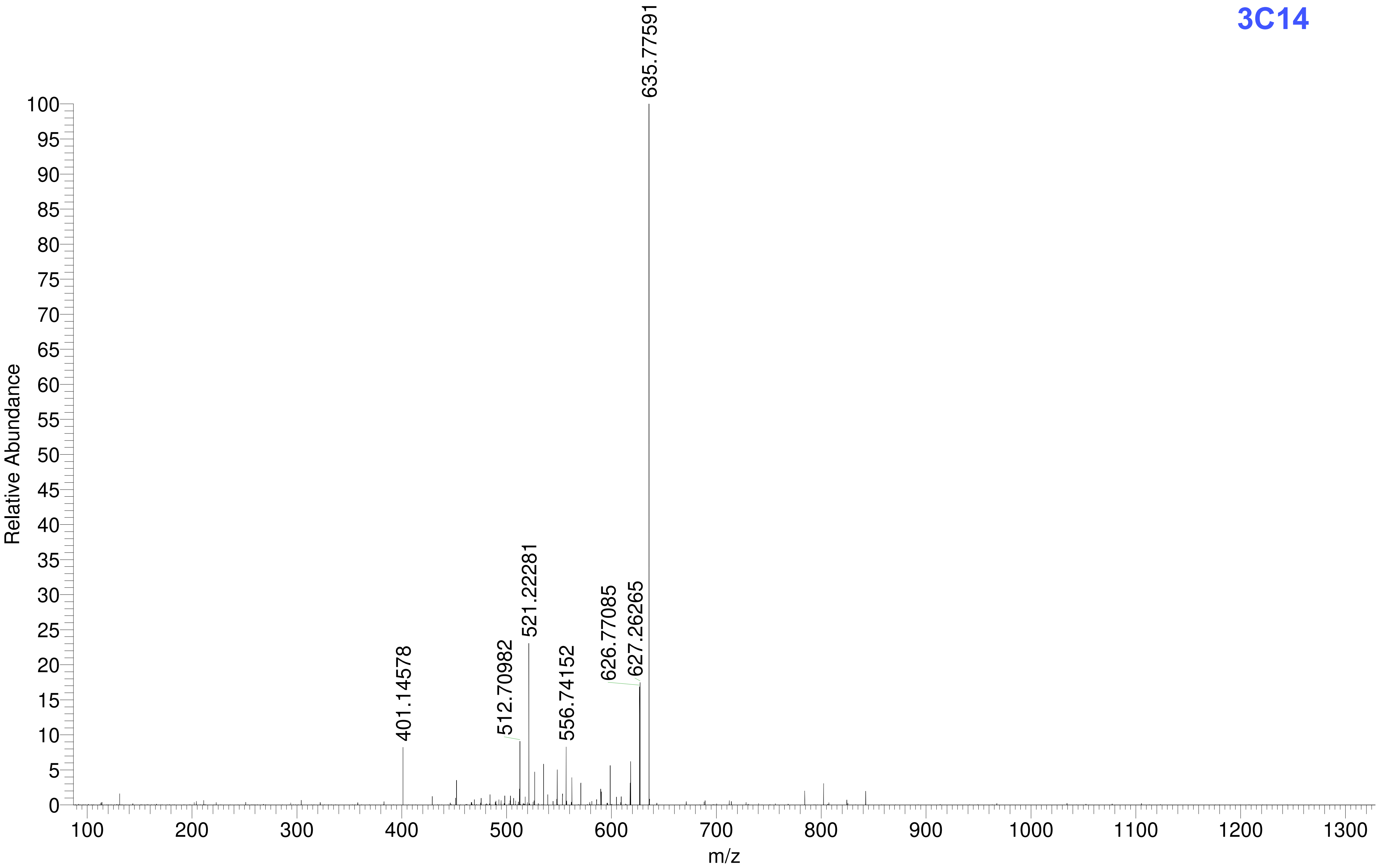
3C14
Relative Abundance
m/z
635.77591
521.22281
401.14578
512.70982
556.74152
626.77085
627.26265
0
5
10
15
20
25
30
35
40
45
50
55
60
65
70
75
80
85
90
95
100
100
200
300
400
500
600
700
800
900
1000
1100
1200
1300

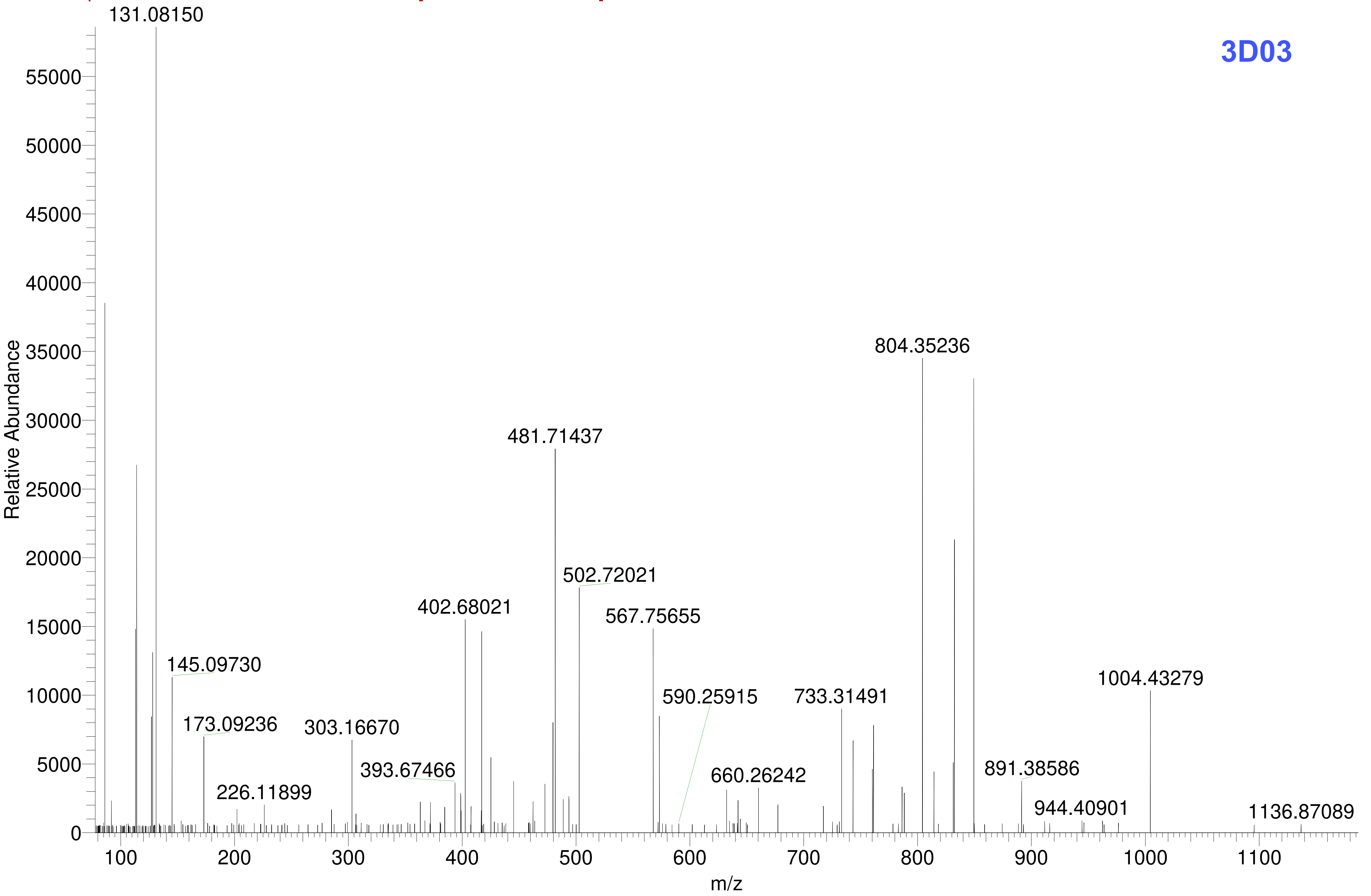
3D03
131.08150
804.35236
481.71437
502.72021
402.68021
567.75655
145.09730
1004.43279
590.25915
733.31491
173.09236
303.16670
393.67466
891.38586
660.26242
226.11899
944.40901
1136.87089
Relative Abundance
m/z
0
5000
10000
15000
20000
25000
30000
35000
40000
45000
50000
55000
100
200
300
400
500
600
700
800
900
1000
1100

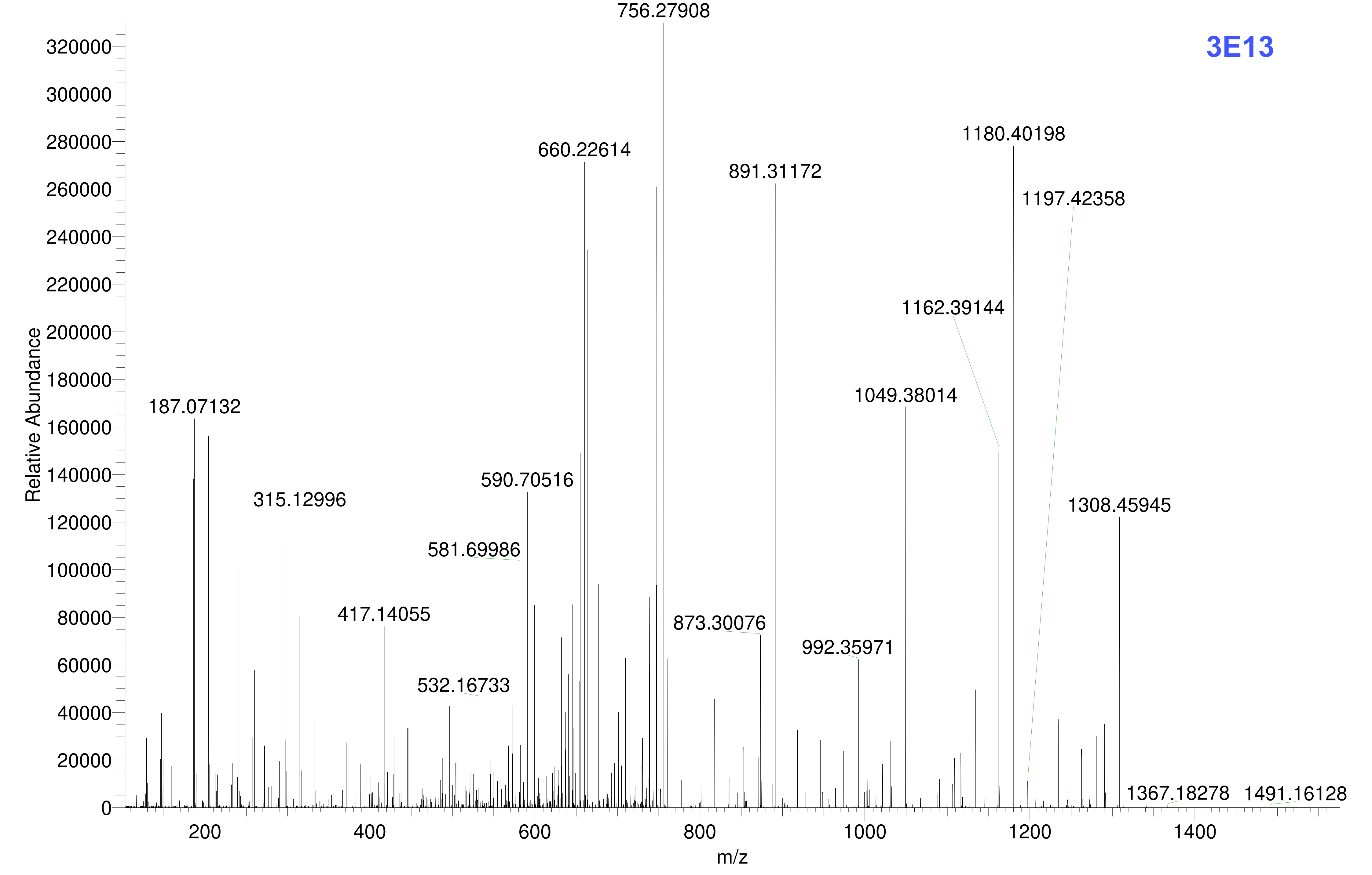
3E13
756.27908
660.22614
891.31172
1180.40198
1197.42358
1162.39144
1049.38014
187.07132
315.12996
590.70516
581.69986
417.14055
873.30076
992.35971
532.16733
1308.45945
1367.18278
1491.16128
Relative Abundance
m/z

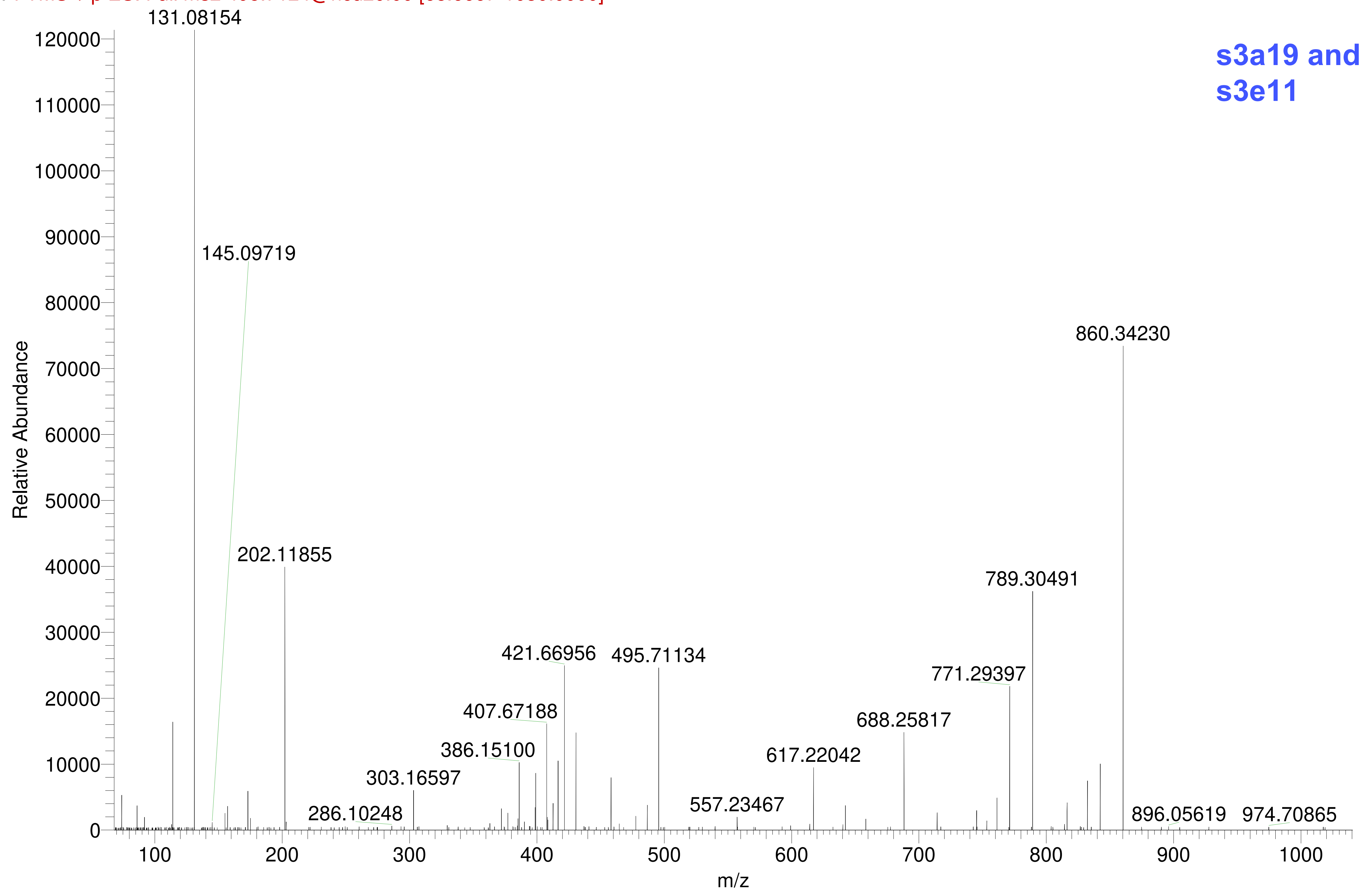

s3a19 and
s3e11
131.08154
145.09719
202.11855
286.10248
303.16597
386.15100
407.67188
421.66956
495.71134
557.23467
617.22042
688.25817
771.29397
789.30491
860.34230
896.05619
974.70865
Relative Abundance
m/z

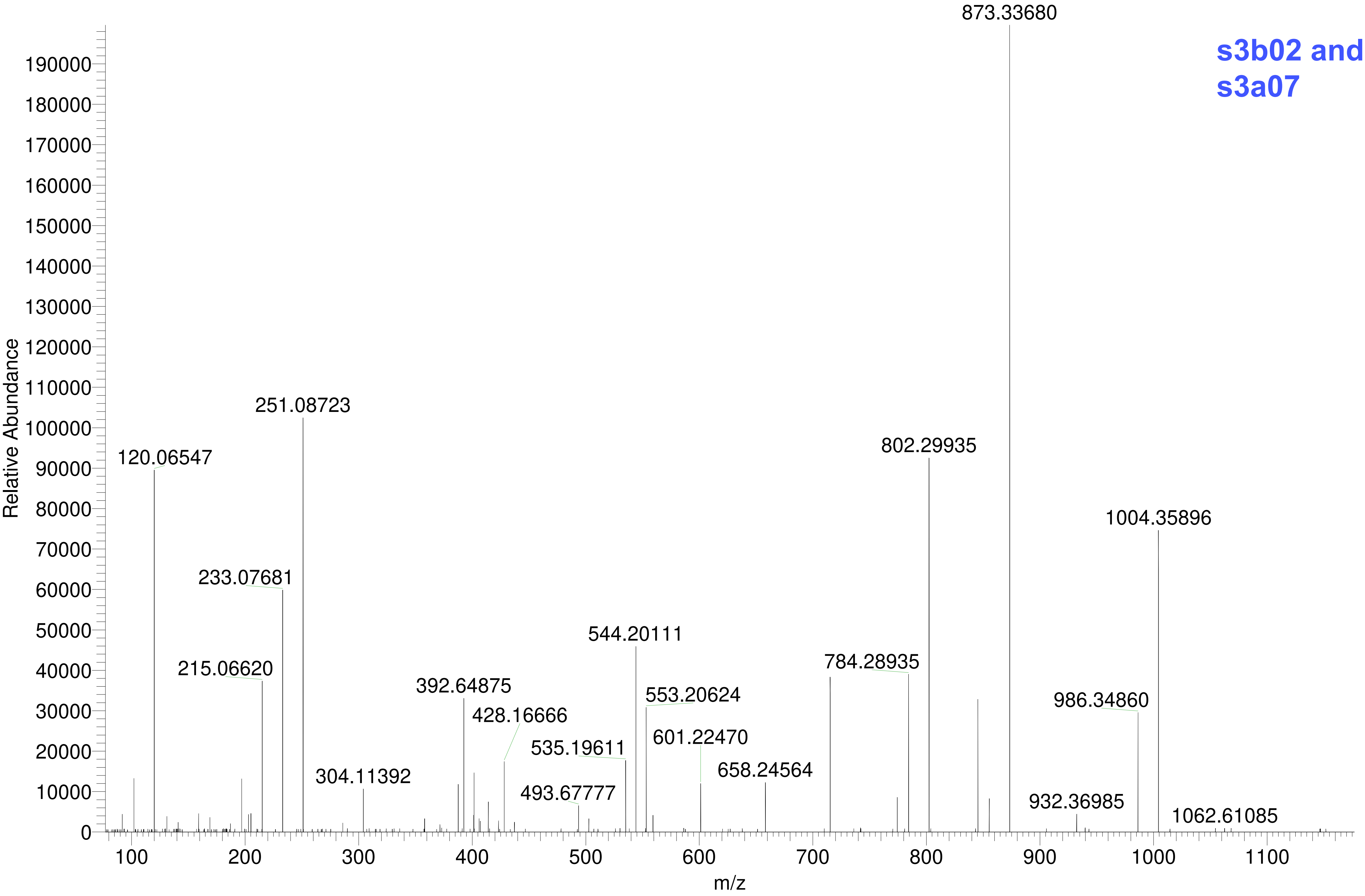

s3b02 and
s3a07
873.33680
251.08723
120.06547
802.29935
1004.35896
233.07681
544.20111
215.06620
784.28935
392.64875
553.20624
986.34860
428.16666
535.19611
601.22470
304.11392
658.24564
493.67777
932.36985
1062.61085
Relative Abundance
m/z

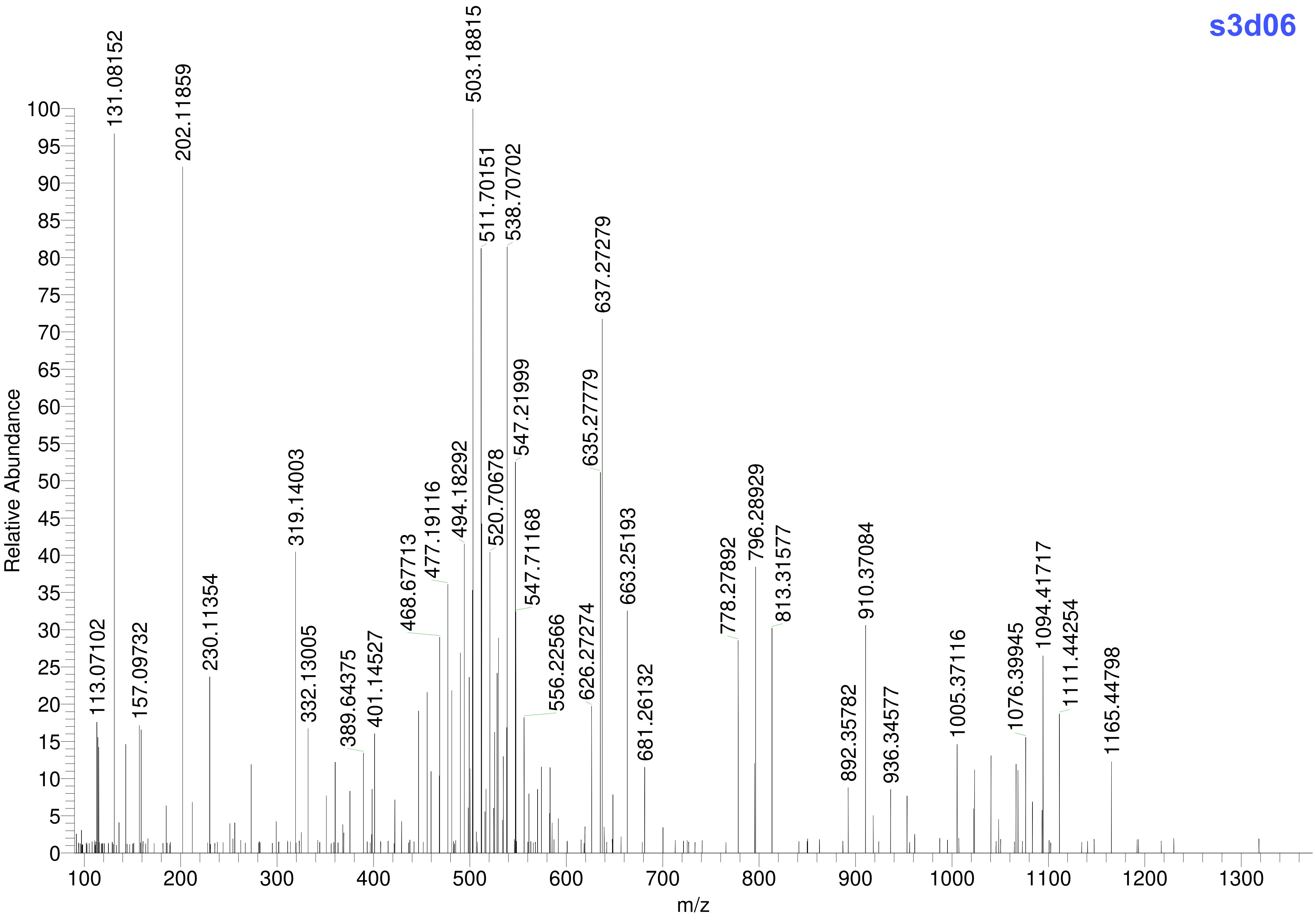
s3d06
Relative Abundance
m/z
131.08152
202.11859
503.18815
511.70151
538.70702
637.27279
635.27779
547.21999
494.18292
520.70678
477.19116
319.14003
547.71168
468.67713
663.25193
796.28929
813.31577
910.37084
778.27892
1094.41717
1111.44254
1076.39945
1005.37116
1165.44798
230.11354
332.13005
401.14527
389.64375
556.22566
626.27274
681.26132
892.35782
936.34577
113.07102
157.09732

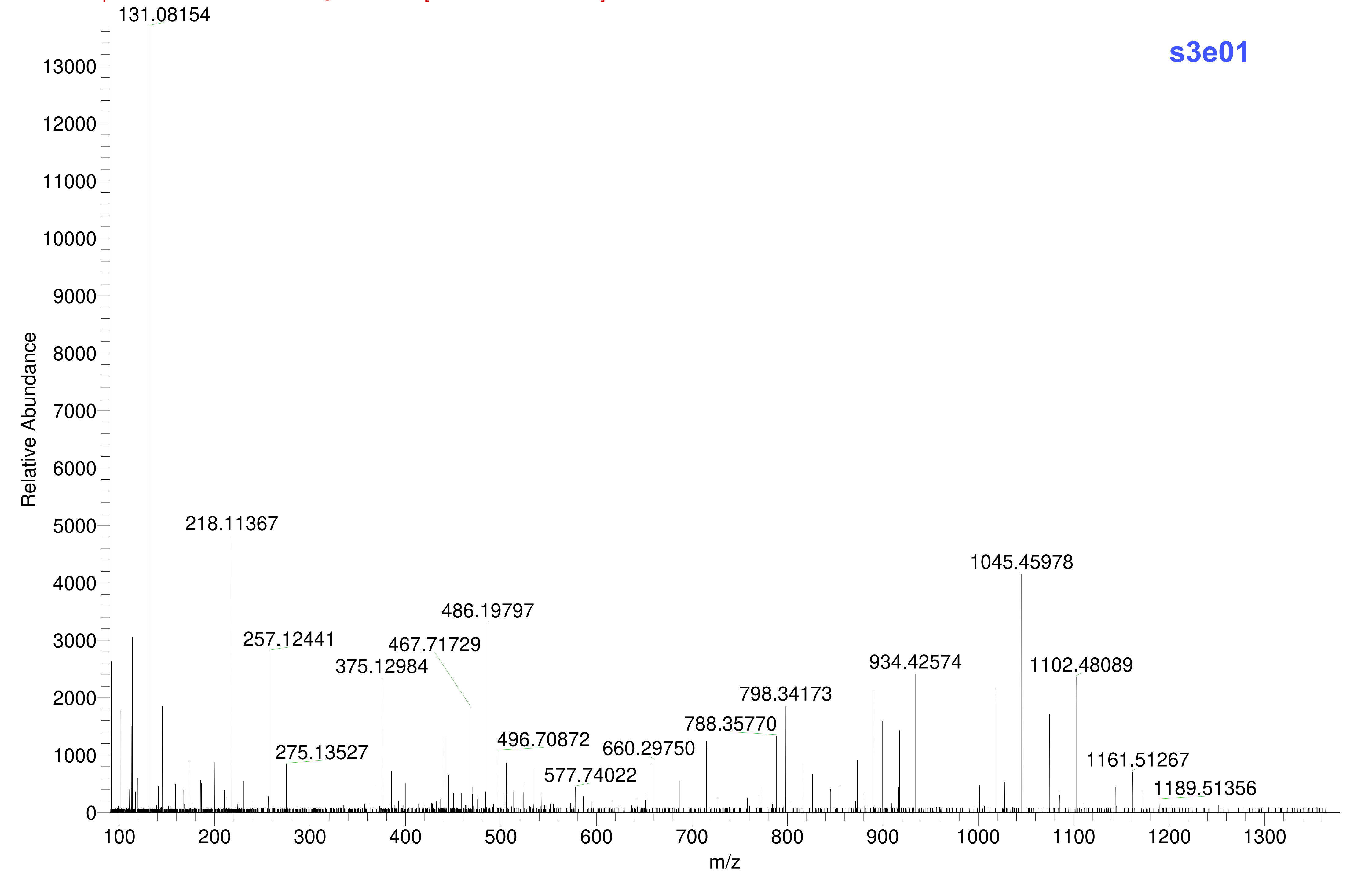
s3e01
131.08154
218.11367
257.12441
275.13527
375.12984
467.71729
486.19797
496.70872
577.74022
660.29750
788.35770
798.34173
934.42574
1045.45978
1102.48089
1161.51267
1189.51356
Relative Abundance
m/z

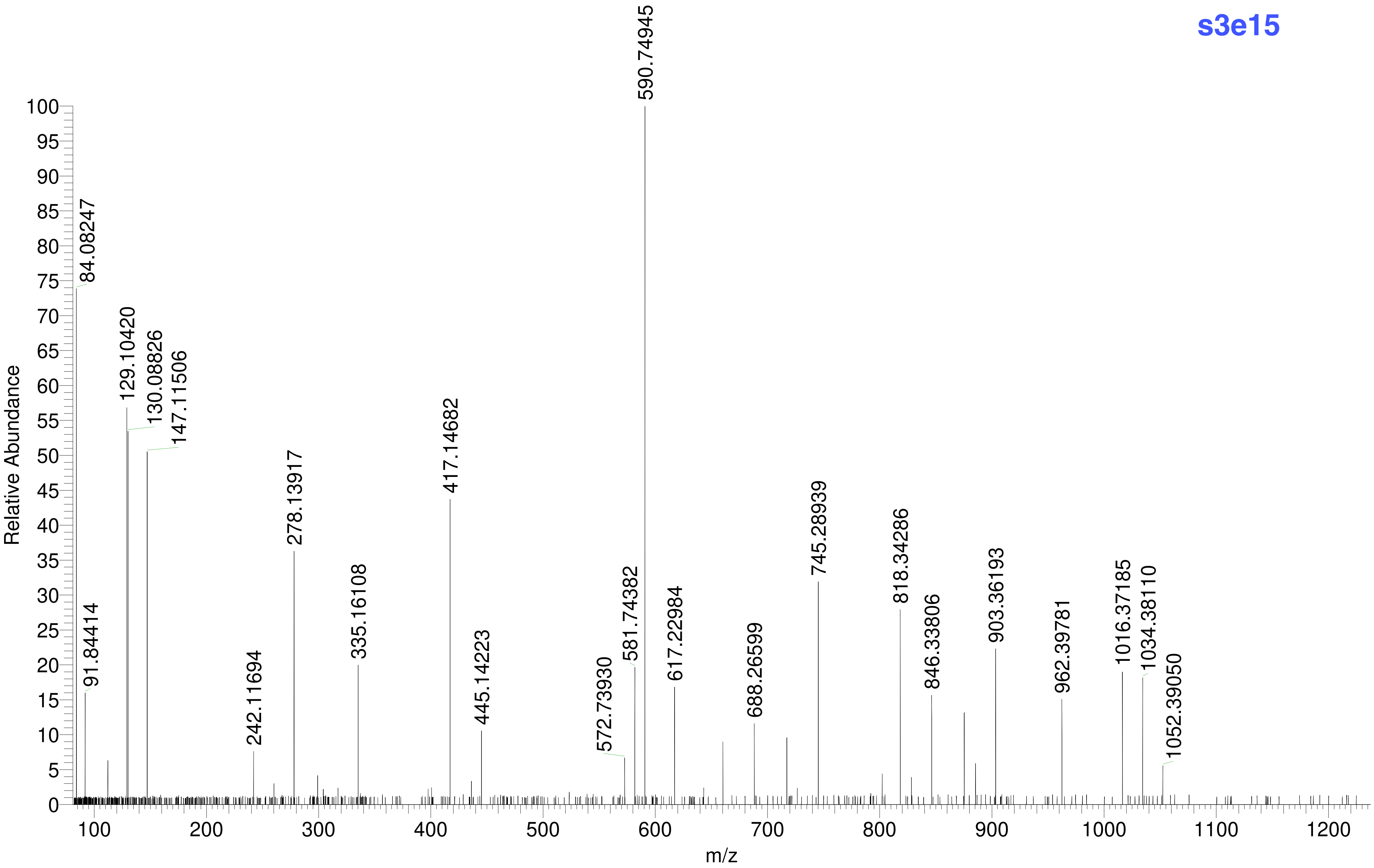
s3e15
Relative Abundance
m/z
590.74945
84.08247
129.10420
130.08826
147.11506
91.84414
242.11694
278.13917
335.16108
417.14682
445.14223
572.73930
581.74382
617.22984
688.26599
745.28939
818.34286
846.33806
903.36193
962.39781
1016.37185
1034.38110
1052.39050
0
5
10
15
20
25
30
35
40
45
50
55
60
65
70
75
80
85
90
95
100
100
200
300
400
500
600
700
800
900
1000
1100
1200
